# The Immobilization of Hyaluronic Acid in 3D Hydrogel Scaffolds Modulates Macrophage Polarization

**DOI:** 10.1101/2024.04.10.588451

**Authors:** Tiah CL Oates, Jasmin Boyd, Louise Dolan, C de Kergariou, Ash Toye, Adam W Perriman, Asme Boussahel

## Abstract

Macrophages are key modulators of immunity, tissue homeostasis and disease development. As our understanding of macrophage biology and their tissue specific behaviors grow the necessity to model macrophages within a 3D biomimetic environment becomes increasingly apparent. Numerous hydrogels have been developed and explored for this purpose, extracellular matrix (ECM) mimicking hydrogels gaining a special interest. In this study, we present the use of such a hydrogel composed of collagen and hyaluronic acid (HA), two of the major ECM components, for the 3D culture of macrophages to model their role in different tissues and diseases. We demonstrate the ability to tailor the mechanical properties of the hydrogel through formulation modulation. Human macrophages retain morphology, viability, and expression of key cell surface markers when 3D cultured within the hydrogel. Interestingly, we demonstrate in this work, that independent of mechanical properties, by adjusting the composition of the hydrogel, specifically HA molecular weight, we can steer macrophage polarization towards either a pro-inflammatory or anti-inflammatory phenotype. This HA-dependent modulation of macrophage behavior is nullified if the HA is chemically crosslinked, shedding light on the impact of one of the most commonly used preparation methods for collagen-HA hydrogels on macrophage behavior.

**Graphical abstract:** 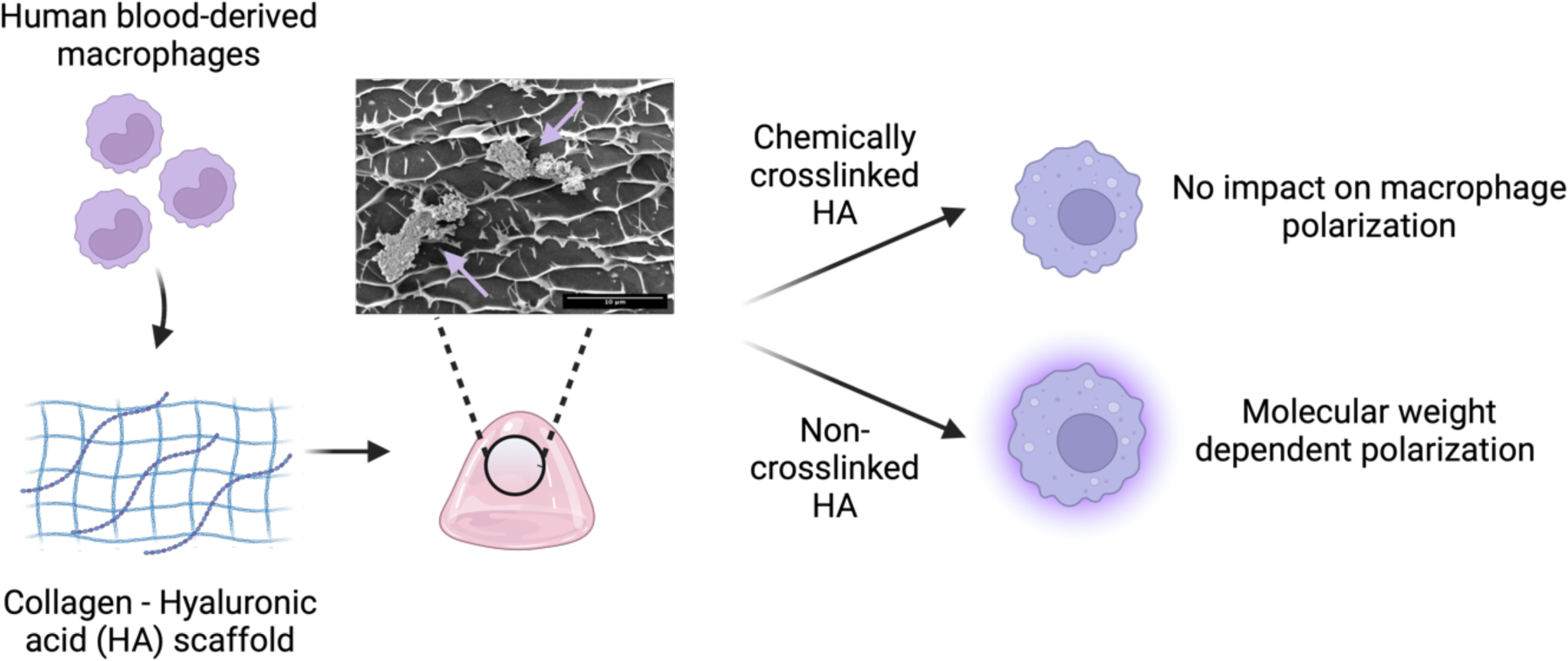

Hyaluronic acid (HA) is an extracellular matrix component, which can modulate the polarization of macrophages. The chemical crosslinking of HA to hydrogel scaffolds counteracts the cell signaling role of this molecule, preventing HA from modulating the immune polarization of macrophages within the model.

**Key points:**

- Collagen-HA hydrogels form suitable biomimetic niches for the 3D culture of human macrophages in which hydrogel composition modulates macrophage phenotype, specifically the presence and molecular weight of HA.
- Polarization of macrophage phenotype by HA is nullified if HA is chemically cross-linked within the hydrogel.

## 1. Introduction

Biomimetic hydrogels allow for the culture of cells within a 3D microenvironment that better recapitulates the extracellular microenvironment of tissues, when compared to 2D cultures. This is achieved by presenting both physical and chemical cues to the resident cells within these hydrogels, resulting in an improved phenotype and function of cells *in vitro*. The development of biomimetic hydrogels is key both in the fields of tissue engineering, and in the *in vitro* study of disease. Ideally, biomimetic hydrogels should emulate the extra cellular matrix (ECM) which is a complex network consisting of a multitude of globular and fibrous proteins that provide structural support for tissues. Collagen, the most abundant protein within the human body, is the main constituent of the ECM. Through fibronectin and integrins collagen interacts with cells facilitating movement through the ECM ^1^. Additionally, collagen acts as an interface between other major components of the ECM including glycosaminoglycans (GAGs) and laminins.

Among the GAGs present within the ECM hyaluronic acid (HA), often referred to as hyaluronan, is of particular interest. Unlike the majority of GAGs, this ubiquitous component of the ECM does not interact with proteins to form proteoglycans, but rather participates in the hydrated network of the ECM along with collagen. The most distinctive property of HA is the ability to bind and absorb significant amounts of water, with the hydration of HA increasing the viscoelastic properties of tissues and tissue elasticity ^2^. Beyond the mechanical properties that HA confers to tissues, this key GAG has been observed to influence both the differentiation and motility of cells ^3,4^.

Macrophages are present throughout the body both embedded in tissues and circulating within peripheral blood, with active roles in immunity and tissue homeostasis. The ability of macrophages to dynamically migrate and respond to external stimuli is essential for the resolution of inflammation within tissues caused by microbial infection or tissue damage ^5,6^. Macrophages dynamically polarize into specialized phenotypes often designated into two subgroups ^7^; the “M1” pro-inflammatory macrophages ^8,9^ and the “M2” anti-inflammatory macrophages ^10,11^, which act in collaboration to perform an array of functions. Critically, beyond this M1/M2 classification a wide spectrum of polarized phenotypes has been noted to exist within macrophage populations ^12^, and macrophages are able to rapidly switch between these phenotypes ^13^.

The ability of macrophages to polarize along this spectrum stems from the sensitivity of these cells to the local tissue environment ^14^. Studies utilizing hydrogels to culture macrophages within a 3D space *in vitro* have highlighted that the composition and stiffness of the microenvironment can impact polarization ^15–17^. Indeed, ECM stiffness has been observed to modulate both the immune signaling pathways through NF-κB ^17^ and integrins ^18^, as well as cell differentiation via the wnt signaling cascade ^19^. The interplay between macrophages and the ECM is reciprocal with macrophages releasing matrix metalloproteinases which remodel the ECM ^20^. This highlights the importance of culturing macrophages in a 3D scaffold that recapitulates the native tissue environment to model their function and behavior *in vivo*.

Further, HA has been noted to stimulate the immunogenicity of macrophages, where macrophages exposed to HA polarize towards different inflammatory states. Within solid tissues free HA normally exists as at a high MW (>1000kDa) ^21,22^, however, the molecular weight of HA within tissues is determined by a range of factors including the presence of hyaluronidases, microbial hyaluronidases, mechanical forces, and oxidization. In 2D *in vitro* macrophages exposed to soluble low molecular weight (MW) HA (<60 kDa) dispersions are induced towards a pro-inflammatory “M1” state, whilst high MW HA (>1000 kDa) suppresses this “M1” phenotype inducing an “M2” anti-inflammatory state ^23–25^. In cases of injury, it has been noted that high MW HA is synthesized contributing to the resolution of the immune response via anti-inflammatory CD44 signalling ^26,27^. Conversely, when HA is catabolized producing low MW HA fragments these bind to TLR4 stimulating a pro-inflammatory response^28^.

This differential response of cells to HA size is a result of HA molecules of different MW influencing the affinity of this protein to cellular receptors ^29^, for example, it has been noted that the binding of low MW HA to CD44 was reversable whereas high MW HA binding was more stable and did not dissociate in the same manner ^30^. Indeed, long polymers of HA have multivalent sites for CD44 biding, whilst smaller molecules have fewer binding sites ^31^. Further, the clustering of receptor complexes by a single HA molecule leads to differential signaling cascades, dependent on the size of the HA molecule ^28,32^. Finally, the size of HA may impact the uptake of the molecule by cells and thus influence the intracellular signaling capabilities of this molecule, such as its association with the receptor for Hyaluronan Mediated Motility (RHAMM) ^33^. Within studies modelling the ECM using 3D hydrogel scaffolds HA is often chemically crosslinked to ensure its retention within the hydrogel ^15,16,34^. However, the impact of immobilized HA within the microstructure of these hydrogels on cells such as macrophages has not yet been fully elucidated.

Here, we have developed a 3D ECM biomimetic model for the study of macrophage immunogenicity. We have shown this collagen-HA hydrogel’s mechanical properties are adaptable through formulation adjustments. It is biocompatible, facilitates the incorporation of migrating human blood-derived monocytes into the system, and supports the differentiation of these cells into mature functional macrophages. Within this 3D environment macrophages are noted to acquire a pro-inflammatory immune profile as seen in previous models ^35^, whilst maintaining expression of anti-inflammatory markers and taking on an intermediate phenotype, a feature of macrophage polarization previously noted by others ^12,36,37^. Utilizing this model system, we investigated both the influence of HA on the mechanical properties of this hydrogel and its role as a cell signalling molecule within the model. This study revealed that soluble HA polarizes macrophages in a molecular weight-dependent manner in 2D culture as shown previously ^21–25^. However, in the hydrogel environment the modulation of macrophage phenotype by HA requires un-crosslinked available HA rather, with the chemical crosslinking of HA in the hydrogel microstructure circumventing this effect.

## 2. Methods

### 2.1. Collagen-hyaluronic acid hydrogel preparation

Bovine collagen solution (Sigma-Aldrich) was diluted to 6.0 mg/ml or 2.5 mg/mL solution (pH 7) in 10X phosphate-buffered saline. The solution was cast into a 24 Well Plate Cell Culture Insert with 8.0 μm Transparent PET Membrane (Falcon) and incubated at 37 °C for 90 minutes. A 0.2 mg/mL hyaluronic acid (HA) solution was prepared using HA sodium salt from *Streptococcus equi* either at 1.5 MDa (∼1.5 - 1.8 MDa) or 50 kDa (∼50 – 70 kDa) reconstituted in 0.1 M MES buffer (pH 5), applied to the top of hydrogels, and incubated at 37 °C for 2 hours and then removed. For crosslinked hydrogels, a 20 mM N-(3-Dimethylaminopropyl)-N’-ethyl carbodiimide hydrochloride (EDC) solution in 0.1 M MES buffer (pH 5), was applied to the top of the hydrogel and incubated at room temperature for two hours to allow the crosslinking of HA to the collagen. Hydrogels were subsequently washed with PBS three times to remove EDC and incubated at 37 °C overnight in cell culture media.

### 2.2. Mechanical testing

The stiffness of the hydrogels was characterized by uniaxial unconfined compressive testing using a Starret FMF5500 single column force testing system (The LS Starrett company), with a 10N load cell and a compressive rate of 1 mm / min displacement rate until just before the incidence of failure. Sample dimensions were measured with an electronic digital calliper (Hika Tools) prior to testing. Both the tangent compressive strength and compressive strain were calculated, converted to positive values, and plotted as a stress-strain graph. Values corresponding to the graph’s linear region (between 10-20 % strain) were selected and a linear fit calculated to obtain the Young’s Modulus, as previously described ^38,39^. The viscoelastic properties of the hydrogels were characterized by rheological testing performed with the Discovery HR 30 (TA instruments) and a 50 mm diameter crosshatched advanced Peltier plate. Strain sweep tested were carried out with measurements between 0.1 % and 100 % strain and a constant frequency of 1 Hz at 25 °C with a gap height of 1000 μm to determine the linear viscoelastic range and the critical strain as previously described ^40,41^

### 2.3. Isolation of human blood monocular cells

Peripheral blood mononuclear cells (PBMCs) were isolated from platelet apheresis blood waste (NHSBT, Bristol, UK) from anonymous healthy donors using experimental protocols approved by Research Ethics Committee (REC 22/WA/0184) using PBMC Lympho Spin (pluriSelect) as previously described ^42,43^. CD14^+^ cells were isolated from PBMCs using CD14^+^ magnetic microbeads (Miltenyi Biotec) as per the manufacturer’s instructions ^37^.

### 2.4. Culture of human macrophages

CD14^+^ cells were cultured at a density of 0.5×10^6^ mL^−1^ in Roswell Park Memorial Institute (RPMI) 1640 (Gibco) medium supplemented with 10 % fetal bovine serum (Gibco), 25 ng mL^−1^ macrophage–colony stimulating factor (w/v; Miltenyi Biotec), and 100U mL^−1^ penicillin/streptomycin (w/v; Sigma Aldrich). Cells were incubated at 37 °C with 5 % CO_2_ for 5 days with full media changes performed every 48 hours. On Day 5 0.02×10^6^ cells were seeded on the top of each hydrogel in 300 μl of media as above, with the addition of 700 μl media to the bottom of the Transwell. For 2D comparisons 0.02×10^6^ cells were seeded into 96 well plates in 300 μl of media. Cells were incubated at 37 °C with 5 % CO_2_ for a further 48 hours before cell characterization assays.

### 2.5. Cryo-SEM microscopy

Hydrogel microstructure was assessed using Cryo-SEM (Quanta 200 - FEI FEG-SEM). Hydrogels were sectioned and mounted with rivets, and A Gatan Alto 2500 Cryo-SEM attachment was used for plunge-freezing, coating, and imaging. Samples were imaged using 16,000x or 4000x magnification. Pore size was quantified using FIJI software (Image J).

### 2.6. Fluorescent Microscopy

Cells in 2D or in hydrogels were fixed in 4 % paraformaldehyde (sigma) for 10 minutes, permeabilized with the addition of 0.1 % Triton X (Sigma) for 15 minutes and incubated with a blocking buffer of 5 % bovine serum albumin in PBS for 1 hour. Cells were subsequently stained with antibodies as per the manufacturer’s instructions for 1 hour. All steps at room temperature with each step followed by PBS washes. The antibodies used were CD14 VioBlue (BD Biosciences, iNOS 594 (Biolegend), CD206 488 (Thermo Scientific), DAPI (Thermo Scientific), and Sytox Green (Invitrogen). Viability was assessed using the LIVE/DEAD™ Cell Imaging Kit (Invitrogen) as per the manufacturer’s instructions. To characterize the incorporation of cells into the hydrogel cells were stained with CellTracker™ Red CMTPX Dye (Invitrogen) prior to addition to hydrogels. Hydrogels were removed from transwells and transferred to glass bottom imaging dishes (Greiner). Confocal microscopy was performed using the SP8 confocal microscope (Leica) using a 63x oil immersion lens. Images were processed using FIJI software (Image J) and fluorescence intensity calculated for individual cells. Distribution of cells within hydrogels analyzed using 3D rendering with LASX software (Leica).

To visualize the distribution of HA within the hydrogels, 0.2 mg/ml of FITC-labelled HA of different molecular weight was added to hydrogels with EDC-crosslinking and incubated in cell media for 48 hours. Hydrogels were removed and imaged by Lightsheet fluorescence microscopy (also known as single plane illumination microscopy) using Z.1 lightsheet system (Zeiss). Images were 3D rendered using Arivis Vision4D software (Zeiss).

To assess macrophage phagocytosis, macrophages were cultured in 2D and in the collagen-HA hydrogel prior of addition of green fluorescent protein (GFP) labelled PODS^®^ crystals (Cell Guidance systems) suspended in macrophage culture media. The GFP-PODS^®^ were incubated with the cells for 24 hours and phagocytosis assessed using confocal microscopy.

### 2.7. Enzyme-linked immunosorbent assay (ELISA)

Supernatant was removed from cells in hydrogels or 2D cultures after 48 hours of culture. The cytokine profile was assessed using the LEGENDplex™ Human M1/M2 Macrophage Panel (Biolegend) as per the manufacturer’s instructions, and samples were acquired using a a MACSQuant flow cytometer (Miltenyi Biotec) and analysed using FlowJo Version 10.7 (BD Biosciences).

### 2.8. HA Leeching

FITC-labelled HA at 50 kDa or 1.5 MDa at 0.2 mg/ml in 0.1 M MES buffer (pH 5), was applied to the top of hydrogels, and incubated at 37 °C for 2 hours. HA was subsequently removed and replaced with 0.1 M MES or 20 mM EDC in 0.1 M MES and incubated at room temperature for 2 hours. Hydrogels were washed with PBS and incubated in cell culture medium at 37 °C. Samples were taken every 24 hours for 96 hours and analyzed using BioTek Synergy Neo2 Reader (Agilent Technologies) with 490 nm excitation and 520 nm emission.

### 2.9. Statistical analysis

Appropriate statistical analyses are detailed in figure legends, along with the number of independent experiments performed. In summary, statistical analyses and generation of graphics were performed using GraphPad Prism 9 (Version 9.1.0) (Dotmatics). Standard deviation is shown where applicable. An unpaired two-sided t-test was used for groups of two sample types. An ordinary one-way ANOVA was followed by a Tukey test to compare every mean to every other mean. Statistical significance is indicated on graphs using standard conventions, as follows: non-significant (ns), p≥0.05; *, p<0.05; **, p<0.01; ***, p<0.001; ****, p<0.0001 *****.

## 3. Results

### 3.1. A collagen-based ECM biomimicking hydrogel with tunable mechanical properties

To facilitate the study of macrophages in immune regulation within tissues an appropriate 3D model is required. A collagen type 1 hydrogel was formed incorporating 200 μg/ml HA as observed *in vivo* in ECM-rich tissues ^44^, and previously utilized in biomimetic hydrogels ^15,45^. To better understand how this collagen-HA hydrogel could recapitulate different tissue ECMs of various mechanical properties^46^, two collagen concentrations (2.5 mg/ml and 6.0 mg/ml) were utilized to regulate hydrogel stiffness. The mechanical properties of this hydrogel were assessed by compression testing (Figure 1a) which showed that increasing collagen concentration from 2.5 mg/ml to 6.0 mg/ml was observed to significantly increase the Youngs compressive modulus of the hydrogel. Further, rheometry measurements showed that increasing the collagen concentration also increased the viscoelasticity of the hydrogel (Figure 1b). It was observed that for both hydrogel compositions the storage modulus (G’) remained greater than the loss modulus (G’’) whilst oscillating strain increased highlighting the elastic properties of the hydrogel in line with reports by Juliar *et al,*. ^47^.

**Figure 1:**
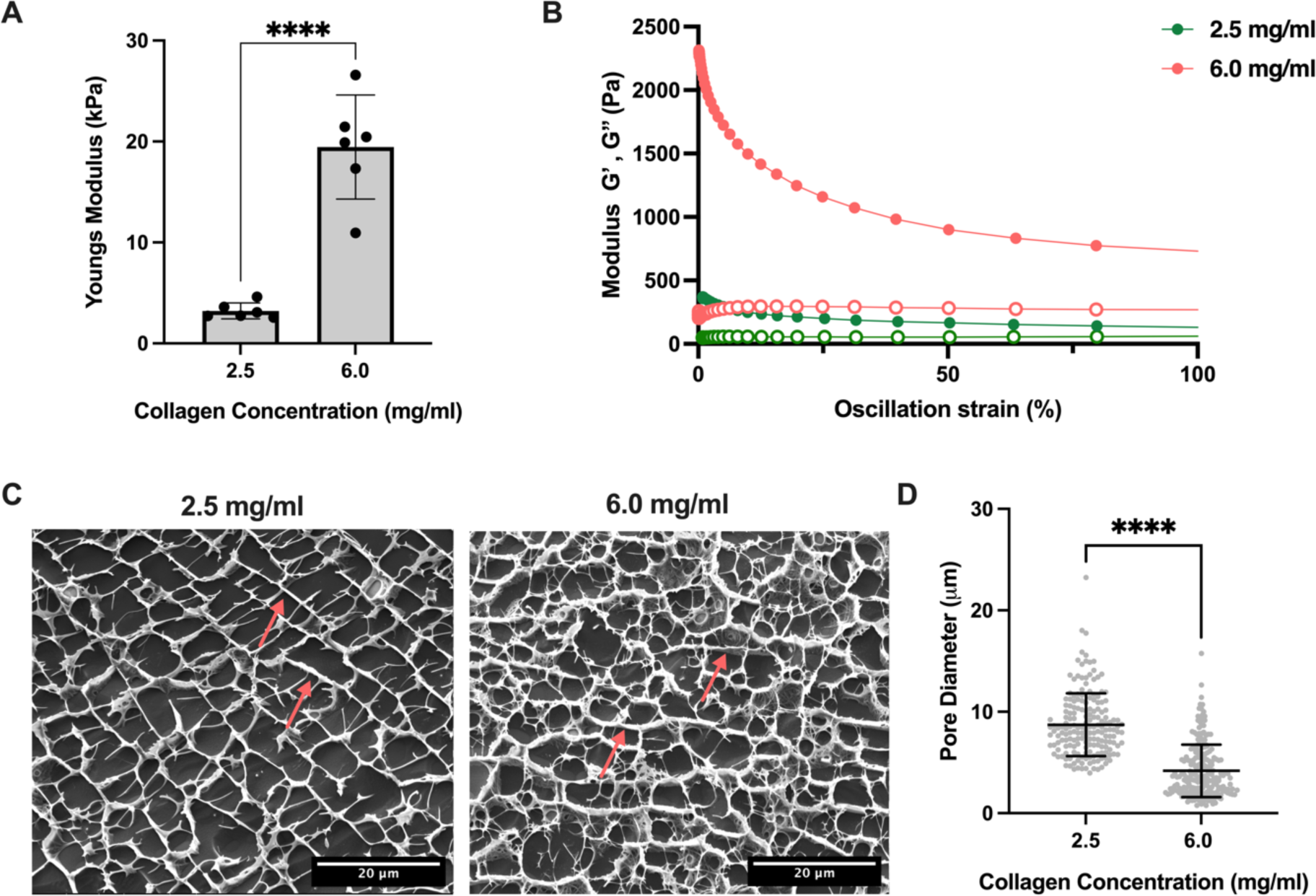
The modulation of mechanical properties in a collagen-based ECM biomimetic hydrogel. a) Youngs modulus of collagen-HA hydrogels, obtained from compression strength testing, at 2.5 mg/ml and 6.0 mg/ml of collagen. Mean and SD shown, (N=3, **** p<0.0001) two-tailed t-test ^47^. b) Rheological characterization of collagen-HA hydrogel at 2.5 mg/ml (green) and 6.0 mg/ml (orange) of collagen illustrating strain dependence in oscillating strain sweeps showing storage modulus G′ (full 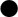) and loss modulus G″ (empty 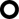). c) Representative cryo-SEM images of collagen-HA hydrogels at 2.5 mg/ml (left) and 6.0 mg/ml (right), scale bar denotes 20 μm, red arrows denote fibres. D) Pore diameter of collagen-HA hydrogel with 2.5 mg/ml and 6.0 mg/ml collagen assessed from cryo-SEM images. Mean and SD shown, (N=3, **** p<0.0001) two-tailed t-test.

To better understand the microstructure and porosity of the collagen-HA hydrogel cryo-scanning electron microscopy (cryo-SEM) was performed, which showed the presence of an organized collagen fiber network within the hydrogels (Figure 1c). A denser microstructure was observed within the 6.0 mg/ml hydrogel when compared to the 2.5 mg/ml (Figure 1c), with a concurrent decrease in pore diameter with increasing collagen concentration (Figure 1d). These changes in morphology at the higher collagen mass fractions, including a reduced pore diameter and increased wall thickness, are consistent with observed increase in the loss and the storage moduli. For instance, at oscillation strains of 1 % and 100 % the Loss and Storage modulus increase by 374.6 % and 469.5 % and 341.2 % and 456.4 %, respectively. The storage modulus increases more than the loss modulus at high and low strains. Consequently, the material becomes more elastic via the increase of collagen mass fraction. 2.5 mg/ml collagen concentration was selected for further experiments, because of the larger pore size and the lower stiffness compared to 6 mg/ml collagen concentration, allowing us to study the influence of other factors on macrophage polarization than mechanical properties.

### 3.2. The culture of human blood monocyte-derived macrophages within the collagen-HA hydrogel

To determine the suitability of the hydrogel scaffold compositions for 3D macrophage culture, the expression of a key macrophage lineage marker CD14 by cells cultured in the hydrogels was compared to standard 2D culture controls (Figure 2a). Human peripheral blood-derived monocytes were differentiated into macrophages and cultured within hydrogels for 48 hours. The expression of CD14 was not significantly altered in macrophages cultured within 2.5 mg/ml or 6.0 mg/ml hydrogels compared to the 2D control (Figure 2b). Similarly, macrophages within both hydrogel compositions maintain viability (>70 %), with no significant difference to those in standard 2D culture (Figure 2c).

**Figure 2:**
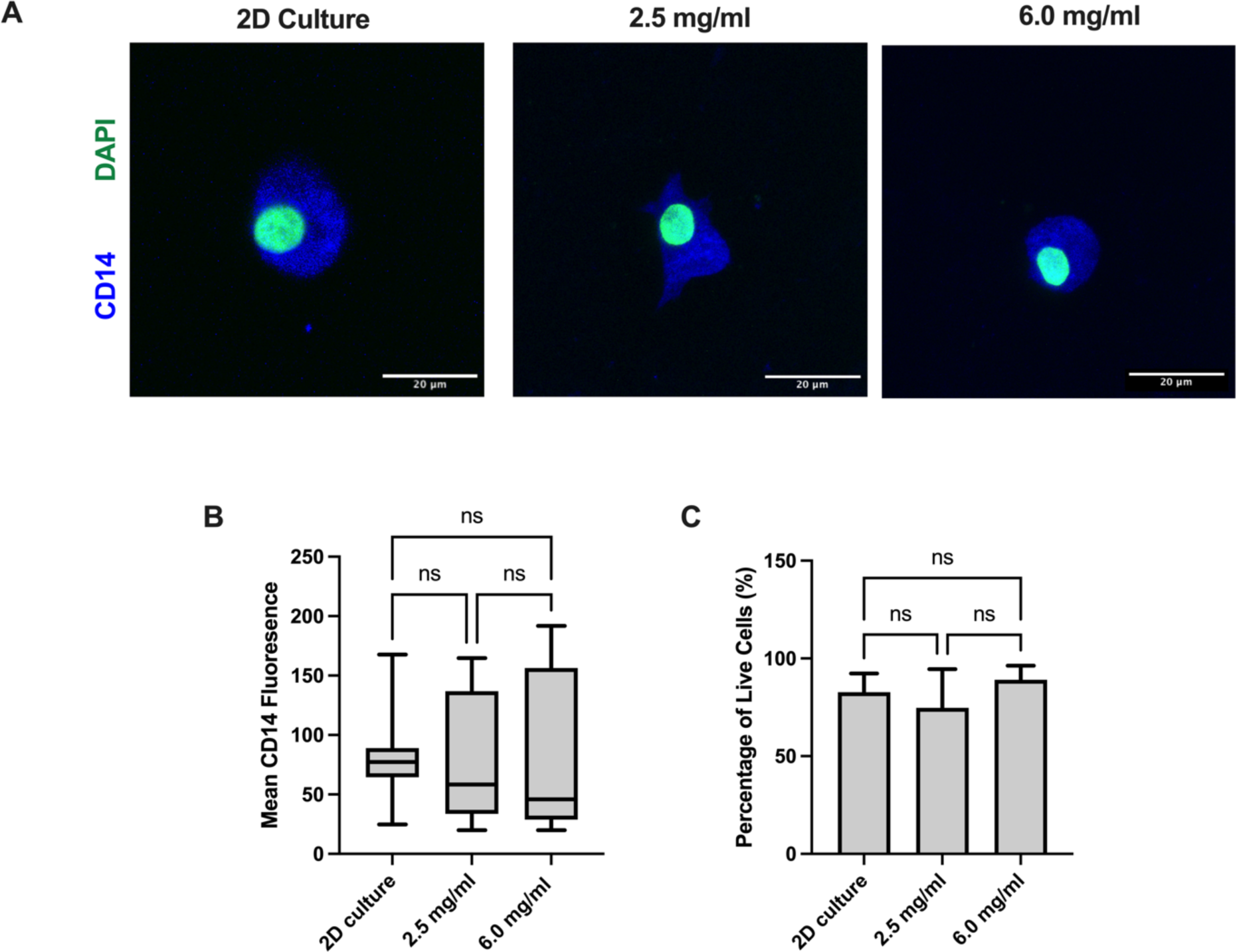
Macrophages retain cell viability and terminal differentiation with hydrogels. a) Representative multi-channel projection overlay of confocal images of macrophages in 2D culture (left) compared to macrophages cultured within 2.5 mg/ml collagen-HA hydrogels (middle) and 6.0 mg/ml collagen-HA hydrogels (right) with CD14 (blue) and DAPI (green) staining, scale bar denotes 20 μm. b) Quantification of CD14 expression of macrophages. Min to max boxplot displaying the mean and SD (N=3 donors, mean 6 cells per donor, ns p≥ 0.05) Ordinary one-way ANOVA with Tukeys multiple comparisons test. c) Viability of macrophages assessed by calcein AM/ ethidium homodimer immunofluorescence. Min to max boxplot displaying the mean and SD, (N=3, ns p≥ 0.05) Ordinary one-way ANOVA with Tukeys multiple comparisons test.

The migration of the macrophages into the hydrogel is key for exploring the immune response of these cells, mimicking the infiltration of macrophages from peripheral blood into the tissue. Lightsheet fluorescence microscopy showed that 24 hours following the addition of CellTracker™ labelled macrophages delivered to the upper surface of the gel (white arrows), they had migrated throughout the scaffold (Figure 3a). Widefield microscopy confirmed that macrophages migrated vertically throughout the entire thickness (2.0 mm) of the hydrogel (Figure 3b). Cryo-SEM highlighted the presence of macrophages within the hydrogel and their size relative to the micropores, establishing that the microstructure supported macrophage motility (Figure 3c).

**Figure 3:**
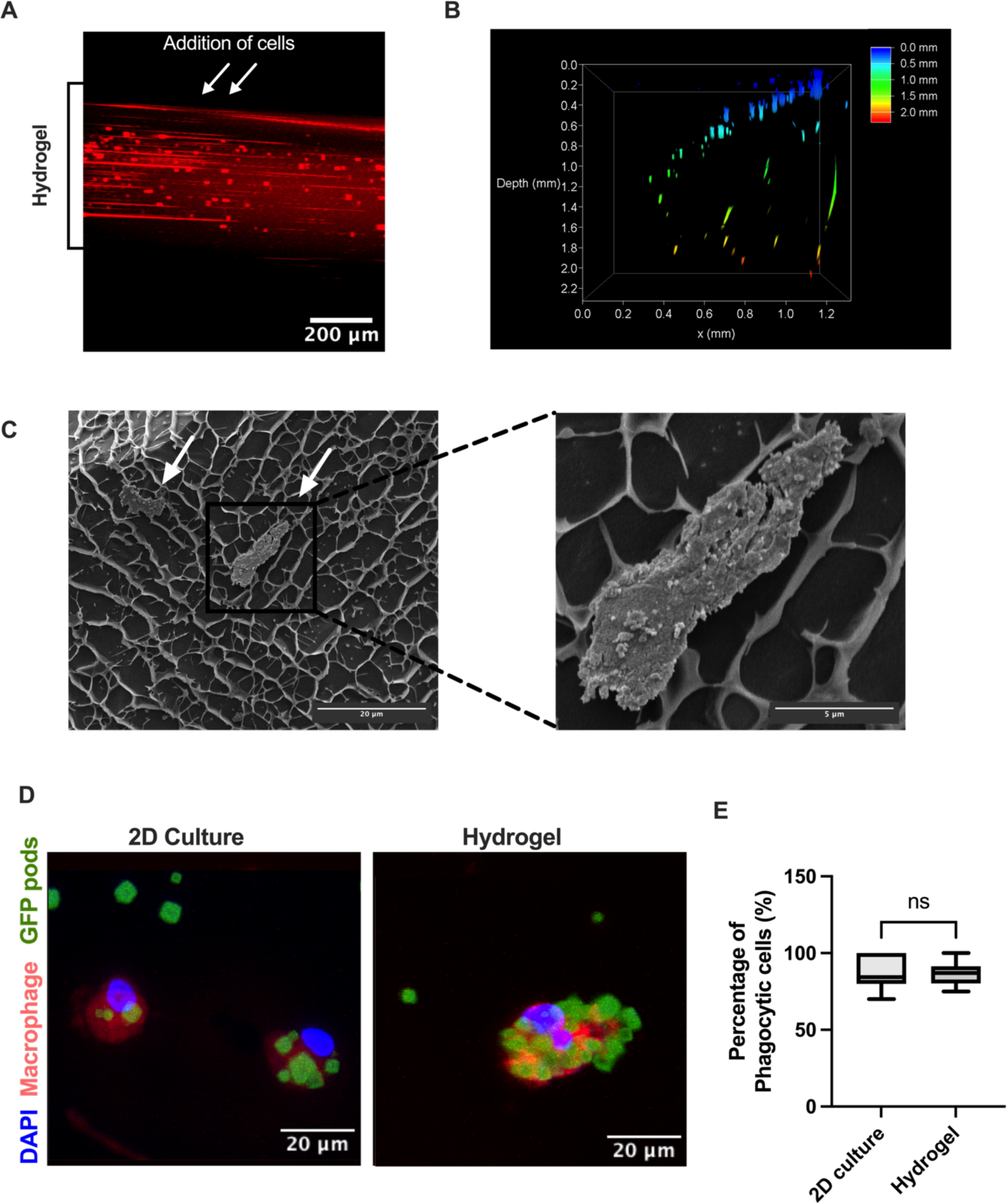
Macrophages are able to perform critical functions within the hydrogel model. a) Representative fluorescent light microscopy image of collagen-HA hydrogel 24 hours post addition of macrophages stained with CellTracker™ Red. White arrows indicate area of macrophage addition, black line indicates width of hydrogel. Scale bar denotes 200 μm. b) Depth of macrophage migration into collagen-HA hydrogel assessed by confocal microscopy. Heatmap scale represents depth where blue denotes 0.0 mm and red denotes 2.0 mm. c) Representative cryo-SEM image of macrophages (white arrows) within hydrogel, at 16,000x scale bar denotes 20 μm (left) and 4000x scale bar denotes 5 μm (right). d) Representative multi-channel projection overlay of confocal images of macrophages in 2D culture (left) compared to macrophages cultured within hydrogels (right) 24 hours post addition of GFP PODS® (green), macrophages labelled with CellTracker™ Red (red) and DAPI (blue), scale bar denotes 20 μm. e) Percentage of macrophages which have phagocytosed at least one of GFP PODS®. Min to max boxplot displaying the mean and SD (N=3 donors, mean 6 cells per donor, ns p≥ 0.05) two-tailed t-test.

The ability of the macrophages to recognize and phagocytose foreign material in 3D was next assessed (Figure 3d). Macrophages within the hydrogel were observed to phagocytose green fluorescent protein (GFP) labelled PODS® at high efficiency following 24 hours incubation Here, with 86.6 % ± 7.3 % of macrophages phagocytosing at least one GFP-PODS®, and no significant difference was noted between the 2D and 3D cultures (Figure 3e).

### 3.3. Macrophages polarize towards a pro-inflammatory M1-like phenotype

It is well established that the inflammatory profile of macrophages is sensitive to the chemical and mechanical stimuli imparted by a hydrogel microenvironment. Accordingly, the polarization of macrophages towards pro-inflammatory or anti-inflammatory profiles within this model was assessed and compared to 2D culture in the presence and absence of inflammatory stimuli. Macrophages within 2D culture stimulated with the pro-inflammatory cytokine IFNγ significantly increased the expression of pro-inflammatory marker inducible nitric oxide synthase (iNOS) (Figure 4a) as expected. Macrophages cultured within the collagen-HA (2.5 mg/ml collagen concentration) hydrogel expressed a pro-inflammatory profile without the addition of IFNγ, significantly increasing expression of iNOS compared to 2D culture (Figure 4a). Surprisingly, expression of the anti-inflammatory marker CD206 was not altered through culture within hydrogels (Figure 4b). Expression of CD206 was, however, significantly decreased in 2D cultures stimulated with IFNγ compared to control macrophages.

**Figure 4:**
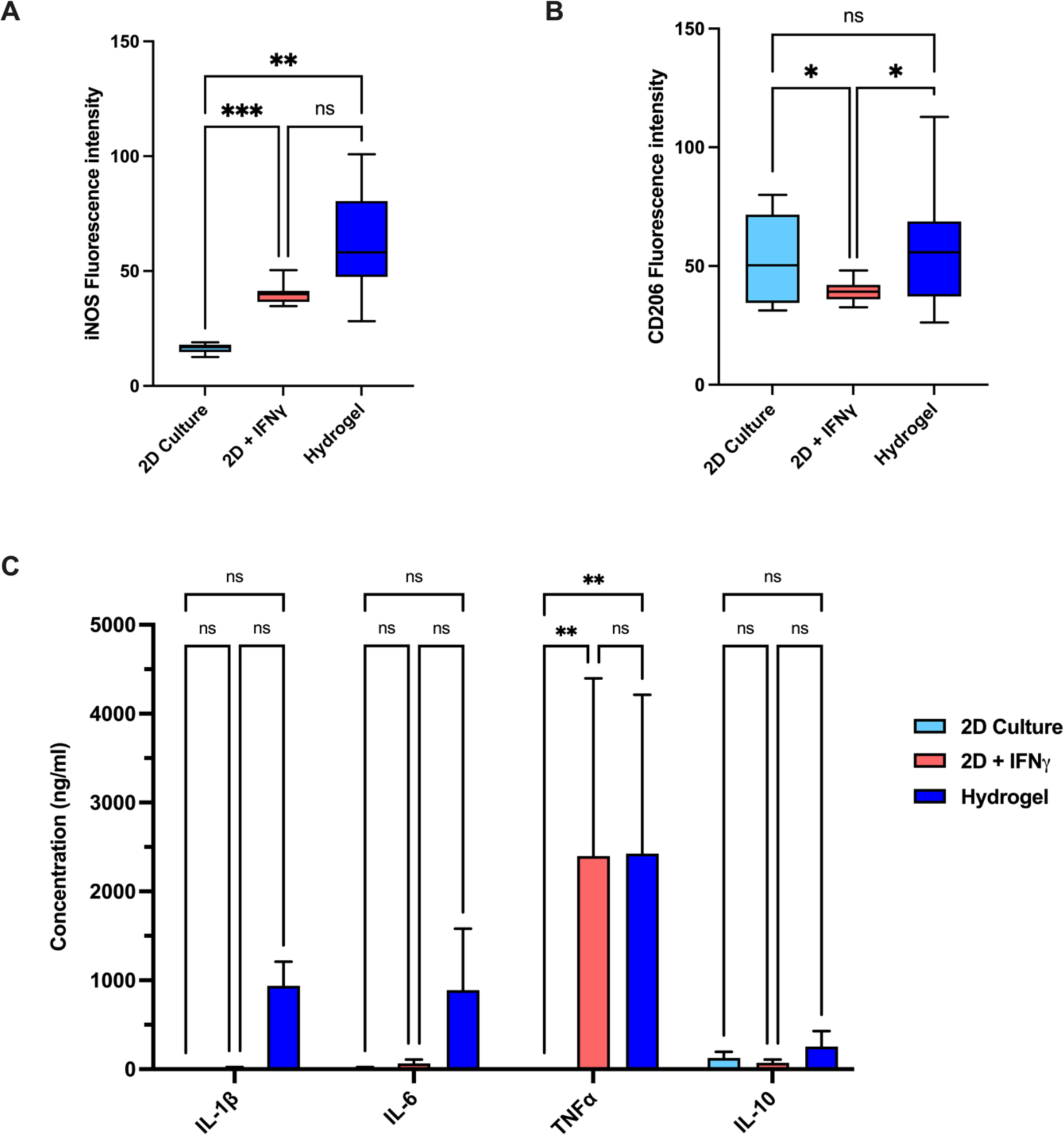
Macrophages polarize towards a pro-inflammatory phenotype within the hydrogel model. a-b) Immunofluorescence quantification showing expression of inflammatory markers of macrophages in untreated 2D culture compared to 2D culture treated with IFNγ for 24 hours, and macrophages cultured within hydrogels for 24 hours. Min to max boxplot displaying the mean and SD, (N=3, **** p<0.0001, *** p<0.001, ** p<0.01, * p<0.05, ns p≥ 0.05) Ordinary one-way ANOVA with Tukeys multiple comparisons test. a) Fluorescence intensity of iNOS. b) Fluorescence intensity of CD206. c) Expression of cytokines measured by ELISA; IL1-β, IL-6, TNFα, and IL-10 in supernatant taken from macrophages in untreated 2D culture compared to 2D culture treated with IFNγ for 24 hours, and macrophages cultured within hydrogels for 24 hours. Mean and SD shown, (N=3, ** p<0.01, ns p≥ 0.05) Ordinary one-way ANOVA with Tukeys multiple comparisons test.

Analysis of secreted cytokines utilizing enzyme-linked immunosorbent assay (ELISA) corroborated this finding (Figure 4c), whereby expression of pro-inflammatory cytokine TNFα was significantly increased in both the IFNγ-stimulated 2D macrophages and hydrogel macrophages compared to the unstimulated 2D control. Similarly, macrophages within the hydrogel expressed the proinflammatory cytokines IL-1β and IL-6, which were not present in 2D cultures. IL-10, a major anti-inflammatory marker, was not significantly altered between the 2D and hydrogel cultures.

### 3.4. HA does not significantly modulate the mechanical properties of collagen-based hydrogels

The biomimetic collagen hydrogel was prepared with 200 μg/ml HA (50 kDa and 1,500 kDa), the incorporation of which was assessed using fluorescein isothiocyanate (FITC)-labelled HA and light sheet microscopy (Figure 5a). This showed high levels of fluorescence penetrating throughout the scaffolds. The mechanical properties that HA imparts on the hydrogel was next assessed. The compressive strength of the hydrogels was not significantly altered by the addition of either 50 kDa or 1,500 kDa HA (Figure 5b). Conversely, as observed previously HA at 1,500 kDa increased the viscoelasticity of the hydrogel ^2^ (Figure 5c). For instance, at oscillation strains of 1 % and 100 % the Loss and Storage modulus increase by 25.0 % and 17.8 % and 21.7 % and 18.3 %, respectively. The loss modulus increases more than the loss modulus at high and low strains. Consequently, the material becomes more viscous via the addition of HA. Cryo-SEM imaging showed an increased presence of smaller microfibrils in hydrogels containing 1,500 kDa HA, suggesting that HA modulates the microstructure by binding to the collagen fibers (Figure 5d). Although, no significant change in overall pore diameter was observed after the addition of HA (Figure 5e).

**Figure 5:**
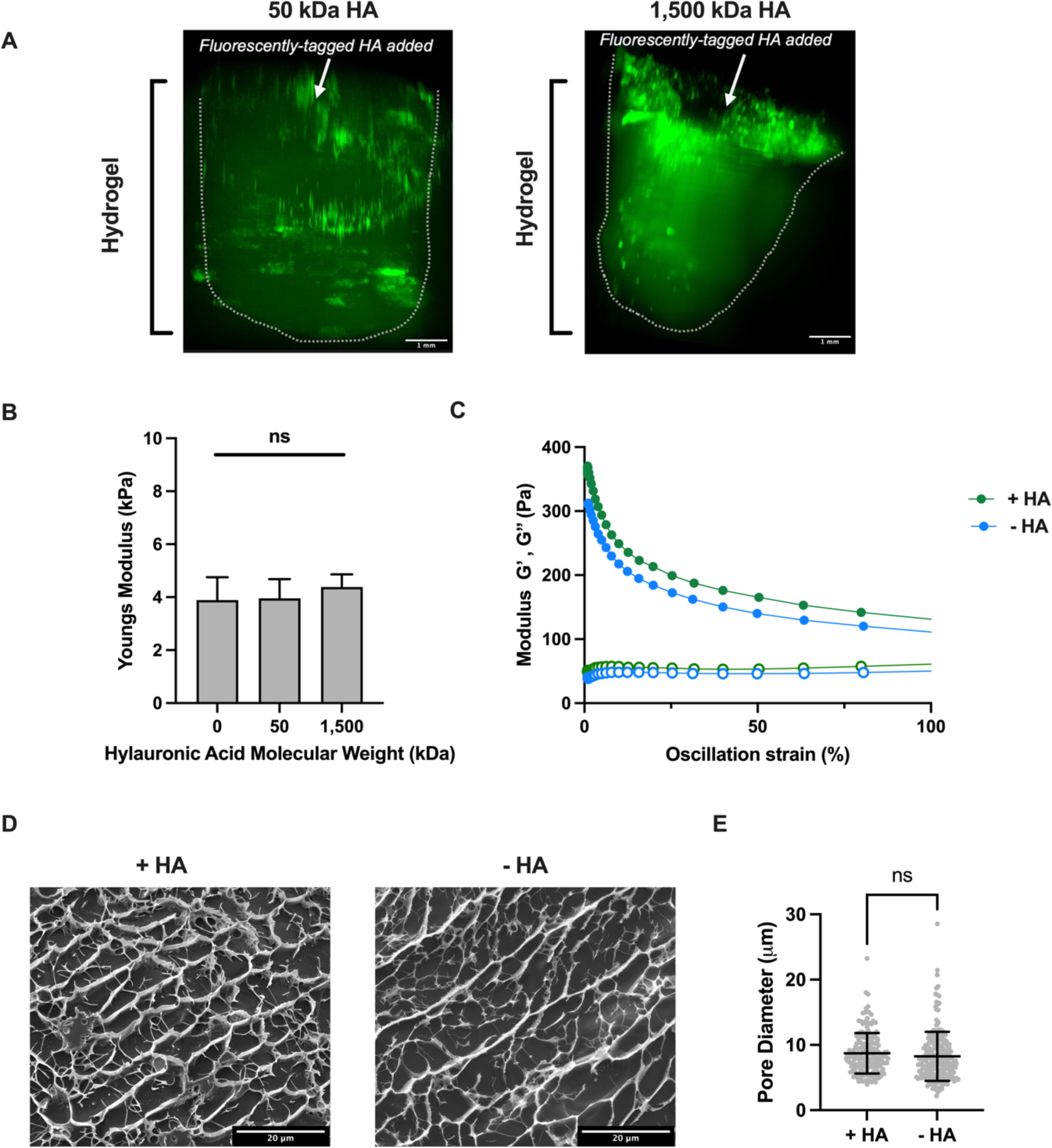
The mechanical properties of collagen-based hydrogels in the presence of HA. a) Fluorescent light microscopy images 24 hours post addition of FITC-labelled HA with a molecular weight of 50 kDa (left) and 1.5 MDa (right) to the collagen-HA hydrogel. White arrow indicates area of FITC-HA addition. Scale bar denotes 1 mm. b) Youngs modulus of collagen hydrogels, obtained from compression strength testing, in the absence of HA compared to 0.2 mg/ml of 50 kDa HA and 1.5 MDa HA. Mean and SD shown, (N=3, ns p≥ 0.05 Ordinary one-way ANOVA with Tukeys multiple comparisons test. c) Rheological characterization of collagen hydrogel in the presence (green) and absence (blue) of 0.2 mg/ml of 1.5 MDa HA illustrating strain dependence in oscillating strain sweeps showing storage modulus G′ (full 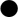) and loss modulus G″ (empty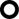). d) Representative cryo-SEM images of collagen hydrogels in the presence (left) and absence (right) of 0.2 mg/ml of 1.5 MDa HA, scale bar denotes 20 μm. e) Pore diameter of collagen hydrogel in the presence and absence of 0.2 mg/ml of 1.5 MDa HA assessed from cryo-SEM images. Mean and SD shown, (N=3, ns p≥ 0.05) two-tailed t-test.

### 3.5. Modulation of macrophage phenotype within hydrogels by hyaluronic acid

To establish the response of the macrophages to HA of different molecular weights, cells were induced with soluble high MW (1.5 MDa) and low MW (50 kDa) HA within 2D culture conditions for 24 hours. As previously observed low MW HA induced a pro-inflammatory phenotype whilst high MW HA induced an anti-inflammatory phenotype ^23^ The expression of iNOS was observed to be significantly increased following treatment with low MW HA, whilst exposure to high MW HA significantly upregulated CD206 expression in 2D macrophages (Figure 6a and b).

**Figure 6:**
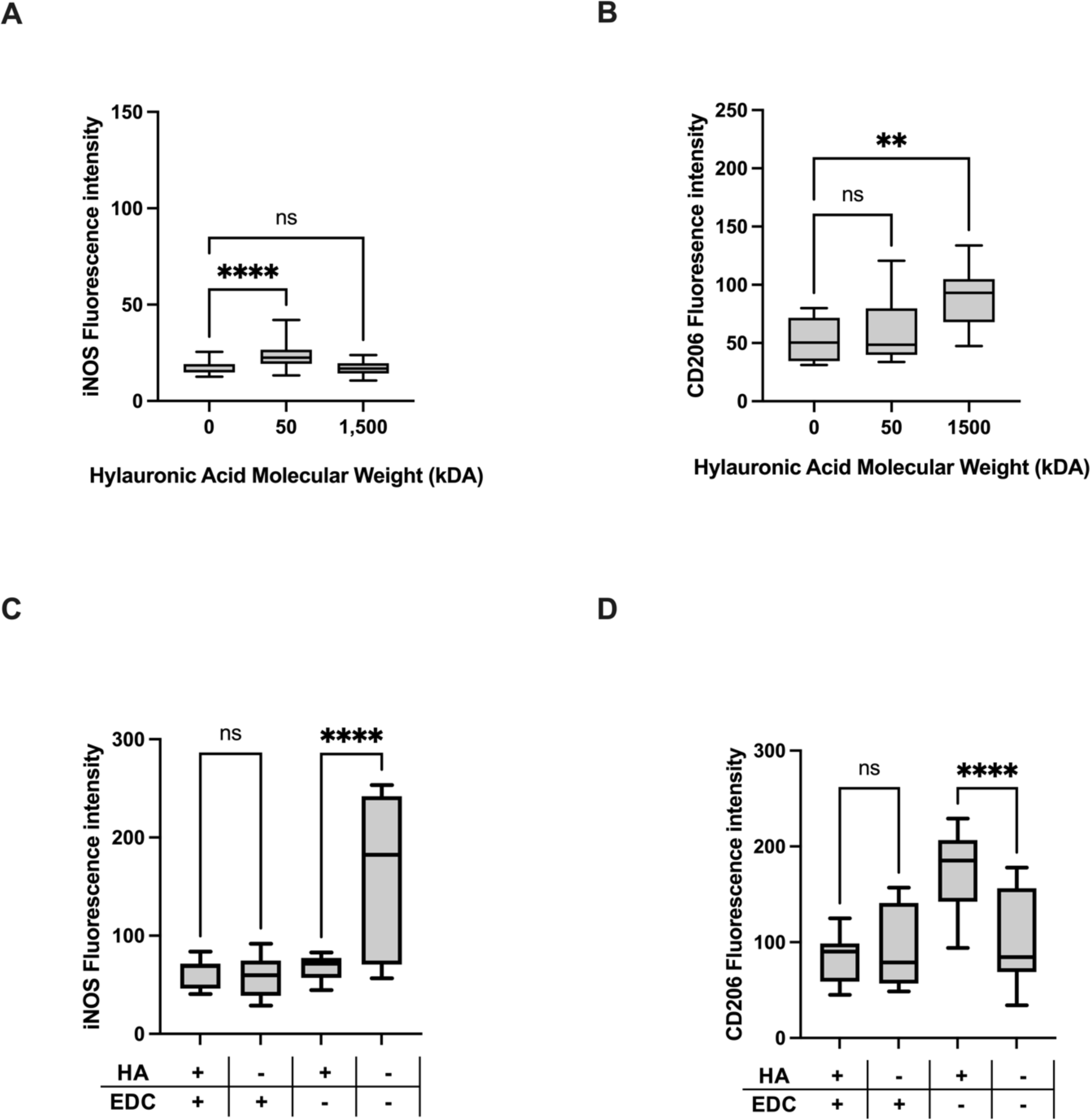
Non-crosslinked hyaluronic acid is required to modulate macrophage phenotype within hydrogels. a-b) Immunofluorescence quantification showing expression of inflammatory markers of macrophages in 2D culture treated with 1,500 kDa or 50 kDa HA for 24 hours compared to untreated control. Min to max boxplot displaying the mean and SD, (N=3, **** p<0.0001, ** p<0.01, ns p≥ 0.05). Ordinary one-way ANOVA with Tukeys multiple comparisons test. a) Fluorescence intensity of iNOS. b) Fluorescence intensity of CD206. c-d) Immunofluorescence quantification showing expression of inflammatory markers of macrophages within collagen hydrogels comprised of 1.5 MDa HA with and without EDC crosslinking, and hydrogels comprised of collagen only, with and without EDC crosslinking. Min to max boxplot displaying the mean and SD, (N=3, **** p<0.0001,, ns p≥ 0.05). Ordinary one-way ANOVA with Tukeys multiple comparisons test. c) Fluorescence intensity of iNOS. d) Fluorescence intensity of CD206.

When incorporated in 3D hydrogels for cell culture, HA is often crosslinked in order to provide structural integrity and increase the mechanical properties of the hydrogel, as well as to prevent leeching of the HA^15,45^. To assess whether the chemical crosslinking of HA within the collagen hydrogel impacts its availability within the model, high MW HA was incorporated into the hydrogel with and without EDC-crosslinking and the polarization of macrophages within the hydrogel assessed. Importantly, no significant difference in the leeching of high MW HA was observed from hydrogels in the absence of EDC crosslinking (Supplemental Figure 1). Macrophages cultured within hydrogels with or without crosslinking were observed to have no significant differences in cell viability (Supplemental Figure 2), indicating that the EDC-crosslinking does not impact the culture of these cells within hydrogels. Significantly, the addition of crosslinked HA had no effect on expression of either iNOS or CD206 compared to macrophages in crosslinked hydrogels containing no HA (Figure 6c and d). However, the expression of iNOS was significantly decreased and CD206 increased in hydrogels containing un-crosslinked high MW HA compared to non-crosslinked hydrogels which did not contain HA (Figure 6c and d). This indicates that macrophages within hydrogel scaffolds polarize in response to available HA, whilst crosslinked HA was observed to have no impact.

## 4. Discussion

Here, we have developed a type 1 collagen-based hydrogel which can be utilized as an ECM biomimetic scaffold for the culture of human cells within a 3D environment. Its tunable mechanical properties allow its adoption for modelling a range of tissues. Increasing collagen concentration of the collagen-HA hydrogel increased both the stiffness and the viscoelastic properties of the model (Figure 1). The mechanical properties of the hydrogel at 2.5 mg/ml collagen were observed to recapitulate the Youngs modulus of *in vivo* dermal strain ^46^ highlighting the potential of the scaffold in modelling key tissues within the body. Notably analysis of the pores created within the collagen-HA scaffold revealed a mean pore diameter of 8.7 ± 3.0 μm in the 2.5 mg/ml collagen-HA hydrogel, ideal for the migration of highly dynamic cells such as macrophages through the material. However, the mean pore size of the 6.0 mg/ml hydrogel was 4.2 ± 3.6 μm and as such is likely to impart strain onto cells, making it more suited to the culture of cells which undergo spreading, such as osteoblasts ^48^. The phenotype of human macrophages was comprehensively characterized within the hydrogel compositions (utilizing a 2.5 mg/ml collagen concentration) in comparison to standard 2D culture. We observe that macrophages retain morphology, viability, and expression of key cell surface markers within the hydrogels. CD14, a pattern recognition receptor fundamental for macrophage innate immunity function was expressed within the hydrogel showing successful differentiation of human primary monocyte-derived macrophages in this system (Figure 2).

The ability of macrophages to migrate in 3D environments is well established and critical to function of these cells ^49^. To establish whether this 3D hydrogel model was suitable for macrophage migration light sheet and widefield microscopy were utilized to observe the vertical migration of these cells into the gel. Indeed, migration was observed throughout the hydrogel in line with work by Li *et al.* where the migration of macrophages into a HA-based hydrogel was observed *in vivo* in the context of soft tissue reconstruction ^50^. Indeed, CryoSEM revealed the integration of macrophages into the microstructure of the scaffold, illustrating the integration of these cells into the hydrogel from the surface. Further, a crucial function of macrophages, undergoing phagocytosis of foreign materials as part of the surveilling immune response ^6^, was supported by the scaffold in which macrophages were able to migrate towards the GFP PODS® and phagocytose them (Figure 3).

The impact of environmental stiffness on macrophage polarization is well established. Indeed, here we corroborate observations by Chen *et al,*. where our collagen-HA hydrogel with a Youngs modulus of 3.22 ± 0.78 kPa induced a pro-inflammatory phenotype in line with the observation that hydrogels with a stiffness of 2.55 ± 0.32 kPa promote an M1 phenotype ^17^ (Figure 4). Increasing the compressive modulus of hydrogels shifts macrophages towards an anti-inflammatory phenotype where hydrogels with a Youngs modulus of 11 and 88 kPa ^51^, 63.53 kPa ^17^, and those with a stiffness of 130 and 240 kPa ^52^, had a reduced capacity to mount an inflammatory response. In hydrogels where the stiffness was further increased to 323 kPa ^17^, and 840 kPa ^52^ macrophages exhibited pro-inflammatory phenotypes, highlighting a scale of stiffness-induced macrophage polarization.

In addition to hydrogel stiffness, the specific components of the hydrogel, and the influence these have on macrophage function and behavior, must also be considered. Indeed, the control of the chemical composition and surface chemistry of the hydrogel are major factors in the modulation of macrophage polarization^53^. As such, here we investigate the role of HA within hydrogels, and the availability of this key signaling molecule, on macrophages. The presence of HA within the hydrogel was confirmed by light sheet microscopy, with FITC-labelled HA observed to be distributed throughout the hydrogel both at low (50kDa) and high (1.5 kDa) MW. Granular regions of higher density were observed, potentially arising from lattice like structures forming within the collagen where pores contained higher concentrations of HA^55^. Further, the leeching of high MW HA from the scaffold was not significantly increased in hydrogels without crosslinking, indicating that HA is present within the scaffold throughout culture.

The presence of HA in the collagen-based hydrogel did not have any impact on stiffness and neither did the molecular weight. Similarly, the pore size was unchanged in the presence or absence of HA, although some microstructural changes are observed in the SEM images (Figure 5). It is thought that HA affects collagen microstructure in a molecular weight dependent manner through electrostatic, hydrophilic and ionic interactions with the collagen fibers. Nevertheless, the outcomes vary depending on the hydrogel formulation, with different studies reporting diverging effects depending on the collagen type, treatment, formulation, and preparation method ^54^. However, HA is expected to affect the viscoelasticity of collagen hydrogels due to its water-drawing capacity. Our observation alight with this, as we noted a discernible impact of HA on the rheological properties of the collagen-HA hydrogel, making it more viscous as previously reported ^2^. In the collagen-HA hydrogel formulation detailed in this study, HA might not exert significant influence over the mechanical properties of the hydrogel, however, we investigated whether the HA composition of the hydrogel might have an impact on cellular behavior possibly through interactions with cell receptors.

We have established the presence of key HA-binding receptors CD44 and TLR4 within macrophage populations illustrating the capability of these cells to respond to HA *in vitro* (Supplemental Figure 3). As seen previously ^23–25^ the molecular weight of HA impacts macrophage polarization in 2D, with low MW (50 kDa) increasing expression of the pro-inflammatory molecule iNOS and high MW (1,500 kDa) increasing expression of the anti-inflammatory marker CD206 (Figure 6). These assays, and those conducted here, utilize soluble HA which is able to interact freely with cell receptors. However, in ECM biomimicking hydrogels HA is commonly crosslinked to provide stability to the scaffold ^15,16,34^. EDC is a commonly used chemical crosslinker, which covalently links HA through the formation of amide bonds, without forming part of the final linkage allowing it to be removed from the scaffold following the reaction ^15,55,56^. Here, we observe that the MW dependent-polarization of macrophages exerted by HA is shown to occur only when HA is available, with the addition of soluble HA to 2D macrophages or in collagen-HA hydrogels in the absence of chemical crosslinking. In hydrogels were EDC-crosslinking was utilized, the addition of high MW HA did not significantly alter either iNOS or CD206 expression. Whereas, in hydrogels in which HA was not covalently crosslinked within the scaffold the addition of this molecule resulted in a reduction in iNOS expression and increase in CD206 expression in line with the impact of soluble HA in 2D.

As such, we suggest that the covalent crosslinking of HA within ECM biomimetic scaffolds sequesters this key signaling molecule and prevents interactions of HA with macrophages. In the body HA interacts with various elements of the ECM including vitronectin, and proteoglycans such as aggrecan integrating HA into the wider macrostructure of tissues, which provides structural integrity within these tissues. Importantly, many of these interactions are reversable and the complex pathways which modulate the crosslinking of HA *in vivo* control the function and availability of this molecule ^57,58^. Further, within the body HA is hydrolyzed by a range of HAases with differential activities and affinities, supporting the bioavailability of HA within tissues. The formation of irreversible amide bonds within the HA molecules in this model therefore does not recapitulate the *in vivo* availability of HA. Crucially, within our hydrogel, although the leeching of low MW HA was significantly higher when EDC crosslinking was not utilized, the leeching of high MW HA was not impacted by EDC crosslinking. This may result from the longer HA chains forming increased electrostatic interactions with collagen and other HA molecules, securing HA within the scaffold.

We present a novel approach for the 3D modelling of macrophage behavior in tissues, providing a comprehensive analysis that considers both the mechanical properties and composition of the hydrogel. We demonstrate the use of an ECM mimetic hydrogel with tunable mechanical properties to support macrophages differentiation, migration, and phagocytosis and to induce a pro-inflammatory macrophage phenotype within the hydrogel. This study showcases the versatility of collagen-HA hydrogels as a platform for the 3D culture of macrophages, allowing the modulation of their behavior, thus offering potential applications in mimicking various tissue microenvironments and pathological conditions. For instance, the mechanical properties of this hydrogel formulation fall within physiological ranges akin to human adipose tissue^46^, together with the resulting macrophage inflammatory phenotype, make this hydrogel system particularly useful in modelling obesity mediated inflammation in the adipose tissue. Notably, our research unravels for the first time the role of chemical crosslinking, a common practice in HA hydrogel fabrication for cell culture for the enhancement of hydrogel mechanical properties and stability, on modulating the biological effect of HA on macrophage behavior. We demonstrate that while uncrosslinked HA within the collagen matrix exerts a molecular weight-dependent influence on macrophage behavior, this effect is nullified upon crosslinking. This insight underscores the intricate interplay between hydrogel composition, mechanical properties, and macrophage behavior, thereby enhancing our understanding of the role of a 3D culture environment on cellular behavior for tissue modelling.

In summary, this study highlights the efficacy of collagen-HA hydrogels as a biomimetic niche for the 3D culture of macrophages and their potential for modelling macrophage behavior in a range of tissues and disease conditions. It also highlights for the first time, the nuanced influence of hydrogel composition, particularly HA crosslinking, on macrophage polarization. This new understanding enhances our ability to replicate macrophage behavior *in vitro* in a more tissue mimetic manner.

## 5. Author contributions

T.C.L.O and A.B conceived and designed the study. A.B secured funding and provided project administration, supervision, and resources. T.C.L.O performed hydrogel synthesis, rheological testing, cell isolation, cell culture, imaging experiments ELISA, flow cytometry, and data analysis. J.B and L.D performed compression testing and preliminary experimentation. C.d.K performed rheological analysis. T.C.L.O and A.B wrote the manuscript. A.T provided flow cytometer resources expertise, secured funding, and provided project supervision. A.W.P secured funding and provided project supervision. All authors contributed to the article and approved the submitted version.

## 6. Acknowledgments

The authors thank Dr Katy Jepson and Dr Judith Mantell of the Wolfson Bioimaging Facility for facilitating the use of the light sheet imaging and Cryo-SEM systems respectively. The authors also acknowledge the Wolfson Foundation and BrisSynBio, a BBSRC/EPSRC-funded Synthetic Biology Research Centre for establishing the Wolfson Bioimaging Facility (University of Bristol) and for the use of microscopes. The authors would like to thank the Bristol Composites Institute for use of Rheology equipment. Schematic representations were generated with the support of Biorender.

