## Supporting information for "The Immobilization of Hyaluronic Acid in 3D Hydrogel Scaffolds Modulates Macrophage Polarization"

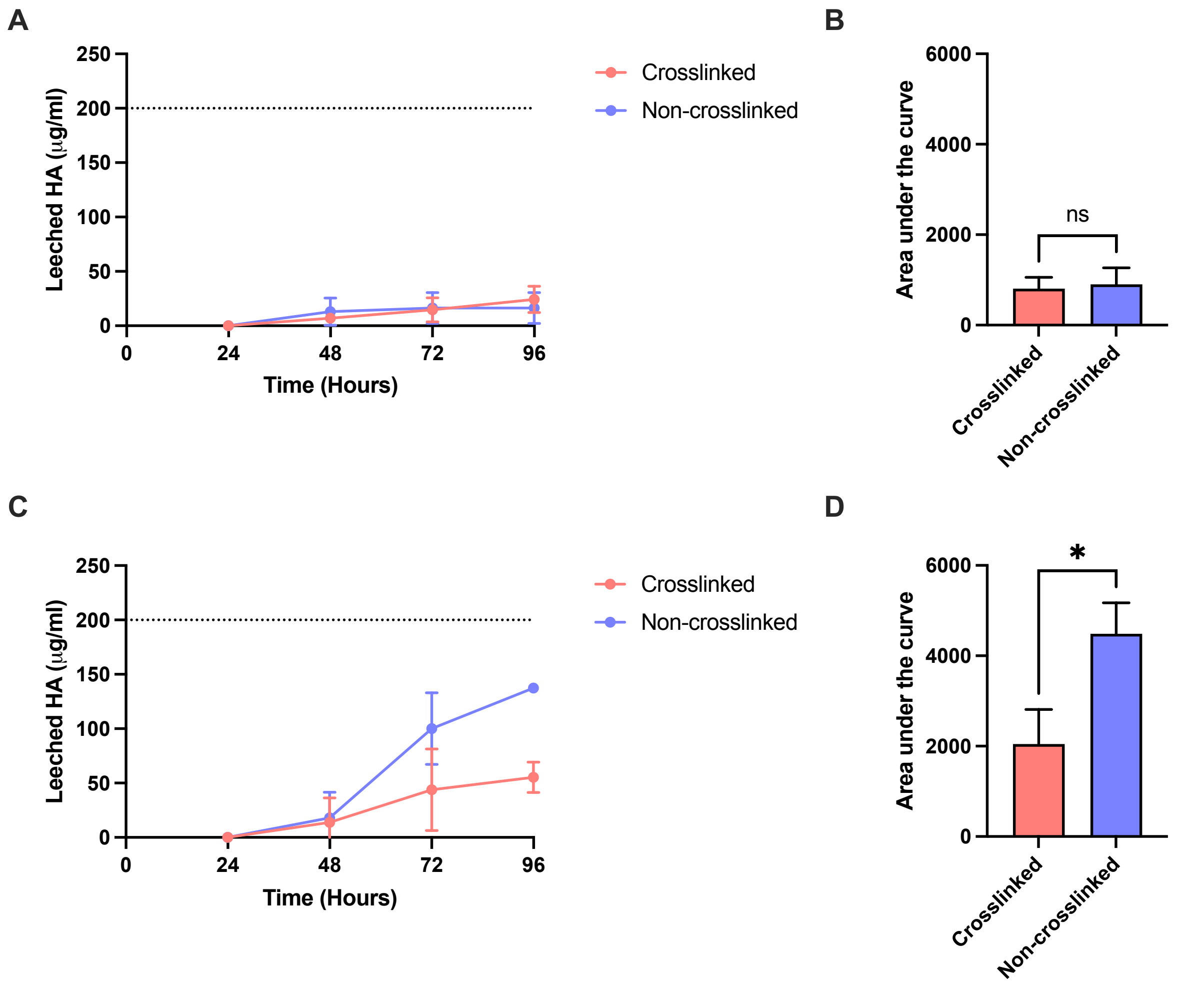


**Supplemental Figure 1: The leeching of HA from 3D collagen hydrogel scaffolds.** a) The mean cumulative concentration of 1.5 MDa fluorescein isothiocyanate (FITC)-labelled HA in media taken from hydrogels at 24, 48, 72, and 96 hours post hydrogel synthesis. Mean and SD shown, (N=3) b) Area under the curve analysis of (a). Mean and SD shown, (N=3, ns $p \geq 0.05$) two-tailed t-test. c) The mean cumulative concentration of 50 kDa fluorescein isothiocyanate (FITC)-labelled HA in media taken from hydrogels at 24, 48, 72, and 96 hours post hydrogel synthesis. Mean and SD shown, (N=3) d) Area under the curve analysis of (c). Mean and SD shown, (N=3, * $p<0.05$) two-tailed t-test.

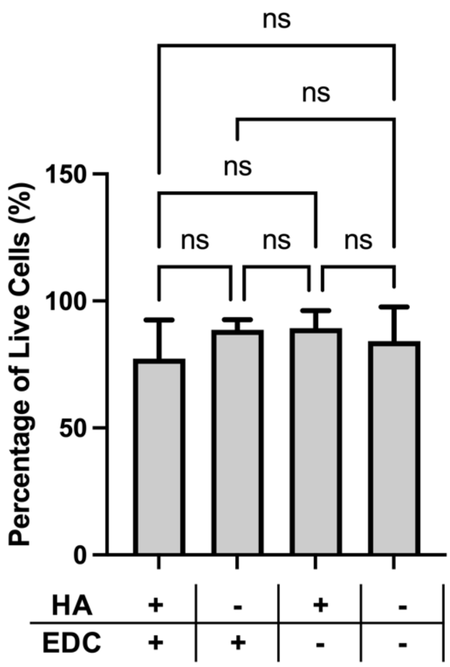


**Supplemental Figure 2: EDC-crosslinking of collagen hydrogels does not impact macrophage viability.** Viability of macrophages assessed by calcein AM/ ethidium homodimer immunofluorescence. Min to max boxplot displaying the mean and SD, (N=3, ns $p \geq 0.05$) Ordinary one-way ANOVA with Tukeys multiple comparisons test.

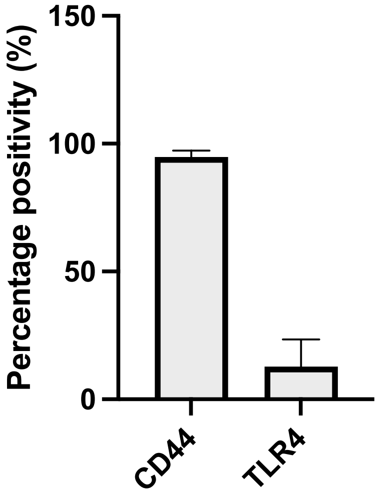


**Supplemental Figure 3: Macrophages express hyaluronic acid receptors CD44 and TLR4.** Percentage positivity expression of CD44 and TLR4 on macrophages as determined using flow cytometry. Min to max boxplot displaying the mean and SD, (N=3).
